# Metabolic scaling has diversified among species, despite an evolutionary constraint within species

**DOI:** 10.1101/2020.05.26.117846

**Authors:** Julian E. Beaman, Daniel Ortiz-Barrientos, Keyne Monro, Matthew D. Hall, Craig R. White

## Abstract

Metabolic rate scales disproportionally with body mass, such that the energetic cost of living is relatively lower in larger organisms. Theory emphasises the importance of fixed physical constraints on metabolic scaling, yet empirical data are lacking with which to assess how evolutionary processes (e.g. mutation, drift, selection) contribute to the observed variation in metabolic scaling across the tree of life. Using a large-scale quantitative genetic study of growth in cockroaches, we show that ontogenetic metabolic scaling is evolutionarily constrained due to an absence of additive genetic variation in juvenile metabolic rate and mass. Using a phylogenetic analysis, we also show that ontogenetic metabolic scaling is more similar among closely related species than among distant relatives, suggesting that the constraints on metabolic scaling are subject to change during lineage diversification. Our results are consistent with growing evidence that there is strong stabilising selection on combinations of mass and metabolic rate within species.

## Introduction

All organisms expend energy to develop, grow, survive and reproduce. The rate at which organisms expend energy, as displayed by their metabolic rate, varies within and among species with consequences for individual fitness within populations (Artacho and Nespolo, 2009, Pettersen et al., 2016) and for the global distribution of matter and energy across ecosystems (Brown et al., 2004; Zaoli et al., 2017). Larger organisms expend more energy overall to sustain a greater mass but, curiously, metabolic rate usually scales disproportionately (allometrically) with body mass such that larger animals have a relatively lower energetic cost of living for their size than smaller animals (Kleiber, 1932, Schmidt-Nielsen, 1984). This allometric relationship is described by a power function where the scaling exponent (or scaling slope on a log-log scale) is positive and less than 1 (White and Kearney, 2014). Metabolic allometry has far reaching implications for macroecological patterns (Zaoli et al., 2017) and ecosystem processes (Allen et al., 2005). Yet the physical and evolutionary processes that govern the origin and maintenance of metabolic allometry remain controversial (West and Brown, 2005, Isaac and Carbone, 2010, White et al., 2019).

Since at least the 1830s, the predominant view on the origin of metabolic allometry has been that there are limitations on metabolic rate arising from the biophysics of organismal structure (Sarrus and Rameaux, 1839, Rubner, 1883, White and Seymour, 2005, Harrison, 2017). For example, geometric constraints on the transport of metabolic inputs and outputs through circulatory networks and across biological surfaces are hypothesised to limit the increase in metabolic rate with increasing body size (West et al., 1997, West et al., 1999, Maino et al., 2014). Systematic variation in metabolic scaling relationships – within and among species (Figure 1) – suggests that the physical constraints on metabolic rate are not immutable (Glazier, 2005, O’Connor et al., 2007, White et al., 2009, Capellini et al., 2010, Uyeda et al., 2017, Norin and Gamperl, 2018). The structural constraints on metabolic rate might be quantitative rather than absolute (*sensu* Mezey and Houle, 2005, Gomulkiewicz and Houle, 2009), allowing individual- and species-level differences in development, physiology and ecology to generate deviations around the average allometric patterns (West and Brown, 2005, Killen et al., 2008, Glazier et al., 2011). For example, phenotypic and environmental sources of variation in metabolic scaling are thought to derive from the energy demands of growth, activity and lifestyle, as well as the effects of temperature, food availability and predation (Glazier, 2010, Killen et al., 2010, Harrison, 2017). Importantly, variation in mass and metabolic rate also has a genetic basis (reviewed in Pettersen et al., 2018), but very little is understood about how microevolutionary processes such as mutation, recombination, inheritance, migration, neutral genetic drift and natural selection contribute to the variation in metabolic allometry across the tree of life (White et al., 2019, Fossen et al., 2019).

**Figure 1.**
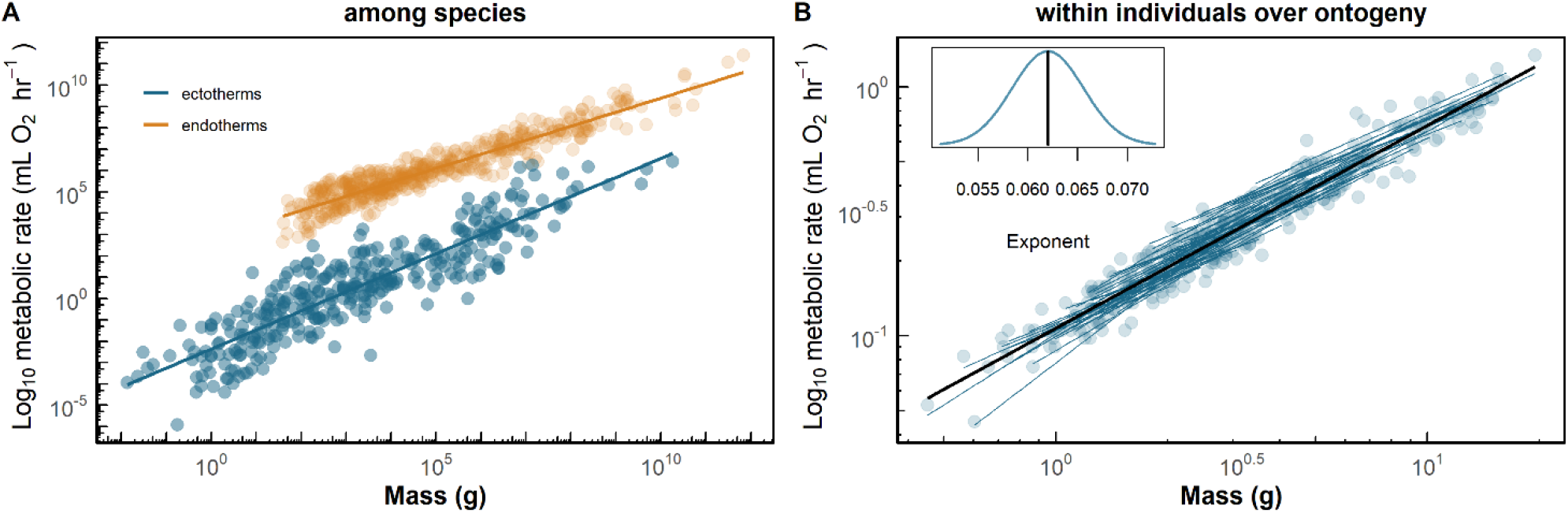
Variation in metabolic allometry at different scales of biological organisation. A: the inter-specific metabolic scaling relationship varies between endothermic and ectothermic taxa. B: the ontogenetic metabolic scaling relationship varies among individual fish (cunner, *Tautogolabrus adspersus*). The inset in B shows the probability density function for the value of the ontogenetic scaling exponent calculated from the mean effect of mass on metabolic rate (0.062, black line) and the standard deviation (0.0036) of individual-level variation in scaling exponents. Data reproduced from publicly available data sets from (A) Uyeda et al. (2017) and (B) Norin and Gamperl (2018).

When the developmental, structural or functional bases of traits are closely shared, such as between mass and metabolic rate, this can generate strong constraints on trait evolution (Blows & Hoffmann 2005). Developmental and functional constraints exert proximate effects on the phenotype and, in doing so, mediate the genetic constraints (Connallon and Hall, 2018) on phenotypic evolution (Arnold 1992). Genetically correlated traits do not evolve independently, which can either restrict or enhance short-term evolutionary responses to phenotypic selection (Lande 1979; Lande & Arnold 1983; Schluter 1996; Agrawal & Stinchcombe 2009). Yet while genetic correlations among traits constrain short-term evolutionary change, at the long timescales over which lineages diverge, the genetic associations among traits can themselves be altered by persistent patterns of directional and stabilising selection (Maynard-Smith et al., 1985; Zeng 1988; Sinervo and Svenson, 2002; Hunt *et al.* 2007). Hence, documenting patterns of genetic variation in the association between mass and metabolic rate is not only a prerequisite for identifying the evolutionary constraints on the metabolic scaling relationship within populations, but will also help to understand how microevolutionary processes contribute to the origin and maintenance of metabolic allometry.

There are now numerous published estimates of additive genetic variance for metabolic rate and mass as separate traits, with heritability estimates for each trait ranging widely from 0 to about ~0.7 (Pettersen et al., 2018). Mass and metabolic rate also usually exhibit a strong and positive genetic correlation (Tieleman et al., 2009, Schimpf et al., 2013, Mathot et al., 2013), and the genetic association between mass and metabolic rate is present in diverse lineages that diverged up to 800 million years ago (White et al., 2019). The widespread and persistent association between mass and metabolic rate suggests that these traits either share many of the same loci (pleiotropy) (Collet et al., 2018) or that the loci underlying mass and metabolic rate are in strong linkage disequilibrium (Slatkin, 2008). No studies to date, however, have investigated whether variation in the mass-scaling of metabolic rate has a heritable genetic basis. This lack of data is a significant impediment to understanding how metabolic scaling evolves within populations in response to age-specific selection on mass and metabolic rate, and for understanding how microevolutionary processes contribute to among-species variation in metabolic scaling relationships.

Here, we focus on ontogenetic metabolic scaling and couple a quantitative genetic analysis of growth and metabolic rate in a cockroach (*Nauphoeta cinerea*) with a phylogenetic analysis of more than 100 species to assess micro- and macroevolutionary patterns of heritable variation in the metabolic scaling relationship. Based on our experimental results and comparative analysis, we suggest that the variation in metabolic scaling is evolutionarily constrained within species, but that these constraints have changed over evolutionary timescales and among lineages. We then propose ways that this hypothesis could be tested experimentally in future research.

## Results and Discussion

### Macroevolutionary variation in ontogenetic metabolic scaling

First, we investigated among-species variation in ontogenetic metabolic scaling relationship using an existing dataset including 133 vertebrate and invertebrate species (Glazier, 2005). The value of the ontogenetic metabolic scaling slope varies considerably among species, ranging between ~0.4 and ~1 (Figure 2). Our analysis (see Materials & Methods) shows that there is a phylogenetic signal in the among-species variation in ontogenetic scaling slope (λ = 0.36 ± 0.13 s.e., LRT statistic = 35.2, df = 1, *P* < 0.0001). In other words, ontogenetic scaling slopes are more similar between closely related species than between distant relatives. Our observation is consistent with there being a heritable basis to the variation in metabolic allometry among species (Lynch, 1991; Hadfield and Nakagawa, 2010); although we note that if closely related species share more similar habitats than distant relatives then this could, in principle, generate an association between phylogenetic relatedness and the metabolic scaling slope.

**Figure 2.**
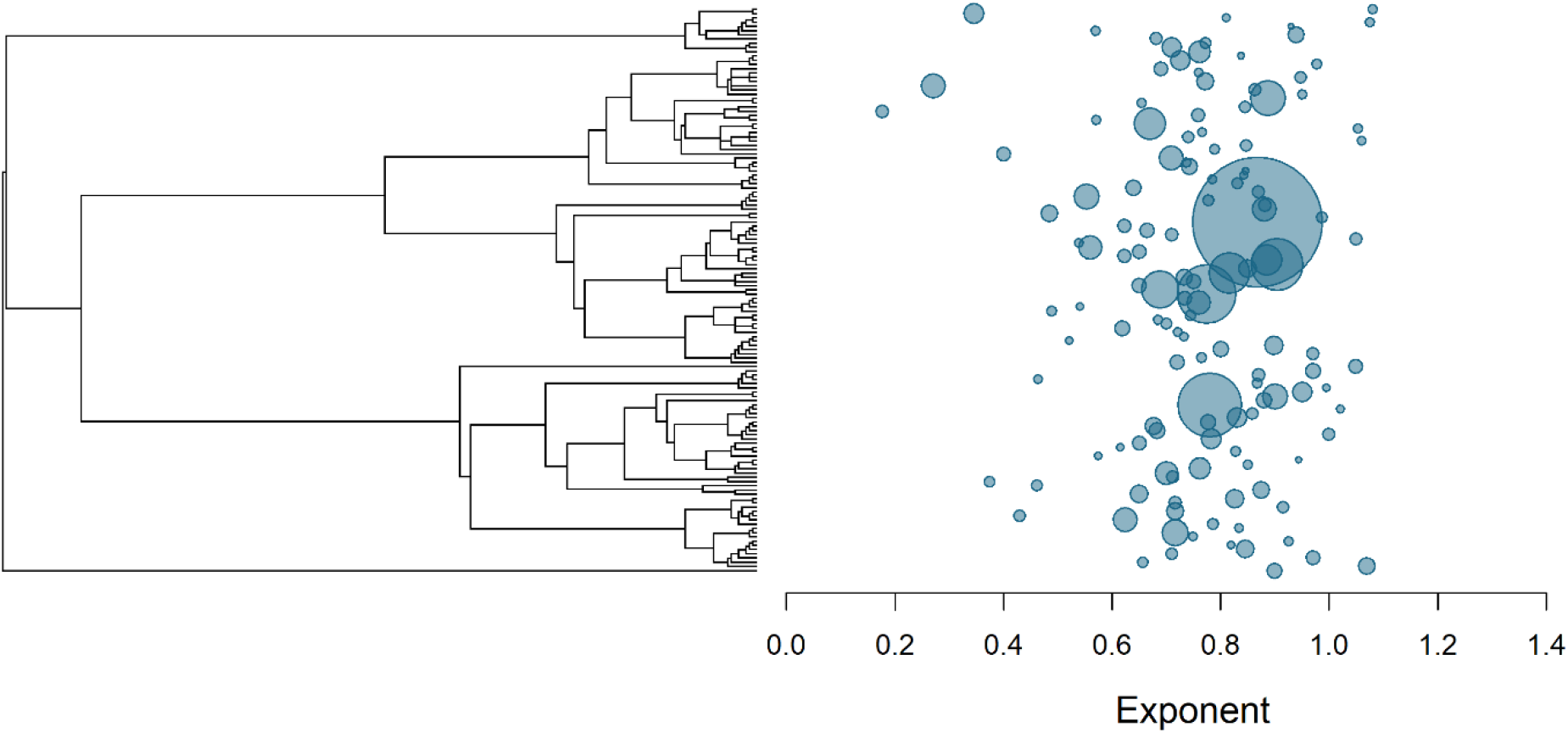
Ontogenetic metabolic scaling slopes exhibit significant phylogenetic variation among vertebrate and invertebrate taxa. More closely related species have more similar ontogenetic scaling slopes than more distantly related species, and there is considerable unexplained variation in the slope once phylogenetic relatedness is accounted for. Point size indicates sample size. Data reproduced from publicly available data sets from Glazier (2005) and reanalysed in the present study.

While accounting for shared evolutionary history, we then tested whether among-species variation in the metabolic scaling relationship was explained by major aspects of animal physiology, development or ecology. We grouped species in the dataset into endotherms or ectotherms, vertebrates or invertebrates, as well as species with aquatic, amphibious or terrestrial lifestyles. We found no consistent differences in the scaling slope between any of the groupings we compared. It is therefore unclear what factors might be driving the among-species variation in the ontogenetic metabolic scaling relationship, but possible sources of variation include variation in energy demanding processes such as growth and activity, as well as environmental effects of temperature (Glazier, 2005; Glazier, 2010; Fossen et al., 2019). In theory, macroevolutionary patterns emerge from microevolutionary processes operating within species (Hansen and Martins, 1996). Recent evidence indicates that the inter-specific metabolic scaling relationship (as opposed to the ontogenetic scaling that we focus on) emerges from patterns of stabilising selection on the covariation in mass and metabolic rate within species (White et al., 2019). There have also been shifts in the inter-specific allometric relationship among major vertebrate taxa, accompanied by relative stasis within clades (Uyeda et al., 2017). Taken together, these studies suggest that the patterns of stabilising selection on the covariation between mass and metabolic rate within species have changed among lineages (Zeng et al., 1988; Hansen and Martins, 1996). Yet there is very little published data with which to assess how the processes of mutation, inheritance and selection operate on combinations of mass and metabolic rate within species and how these processes differ between species. We acknowledge that macroevolutionary variation in metabolic scaling might also arise due to higher-level processes, such as species selection, but theory on multilevel selection is still being developed (Chevin, 2016) and here we restrict our focus to microevolutionary processes.

### Microevolutionary constraints on ontogenetic metabolic scaling

Evolutionary responses to selection require there to be a heritable genetic basis to trait variation (Walsh and Lynch, 2018). No studies to date have obtained the data required to rigorously assess how much of the variation in metabolic scaling relationships is due to heritable additive genetic effects while also accounting for other known sources of variation such as maternal effects, which can confound estimates of heritability (Hadfield and Kruuk, 2007). We estimated the additive genetic variance in the ontogenetic scaling slope and intercept from a large sample of speckled cockroaches (*Nauphoeta cinerea*) of known pedigree. To do so, we measured metabolic rate (2687 measurements) repeatedly in individuals (1269 individuals with 2 – 7 repeated measures each) as they grew in size during development (Materials and Methods). This represents one of the largest sampling efforts ever undertaken for a physiological trait like metabolic rate. Our data show that the metabolic scaling slope and intercept are not fixed quantities and displayed phenotypic variability arising from maternal effects and other unaccounted for sources of variation (Table 1, Figure 3). Yet we detected little additive genetic variance in the metabolic scaling slope and intercept, despite having the statistical power to detect additive genetic variance in growth rate and adult mass (Table 1). The presence of maternally derived variation in the way that metabolic rate increases with mass during development provides the opportunity for selection on different combinations of age- and mass-specific metabolic rate. While we did not estimate selection gradients on mass and metabolic rate in our study, even if selection favoured particular combinations of mass and metabolic rate across development, the mean scaling relationship would not be expected to show an evolutionary response to selection due to a lack of heritable genetic variation underlying the mass-scaling of metabolic rate. The major implication is that while the ontogenetic metabolic scaling relationship is not phenotypically fixed, it does appear to be evolutionarily constrained within species.

**Table 1.**
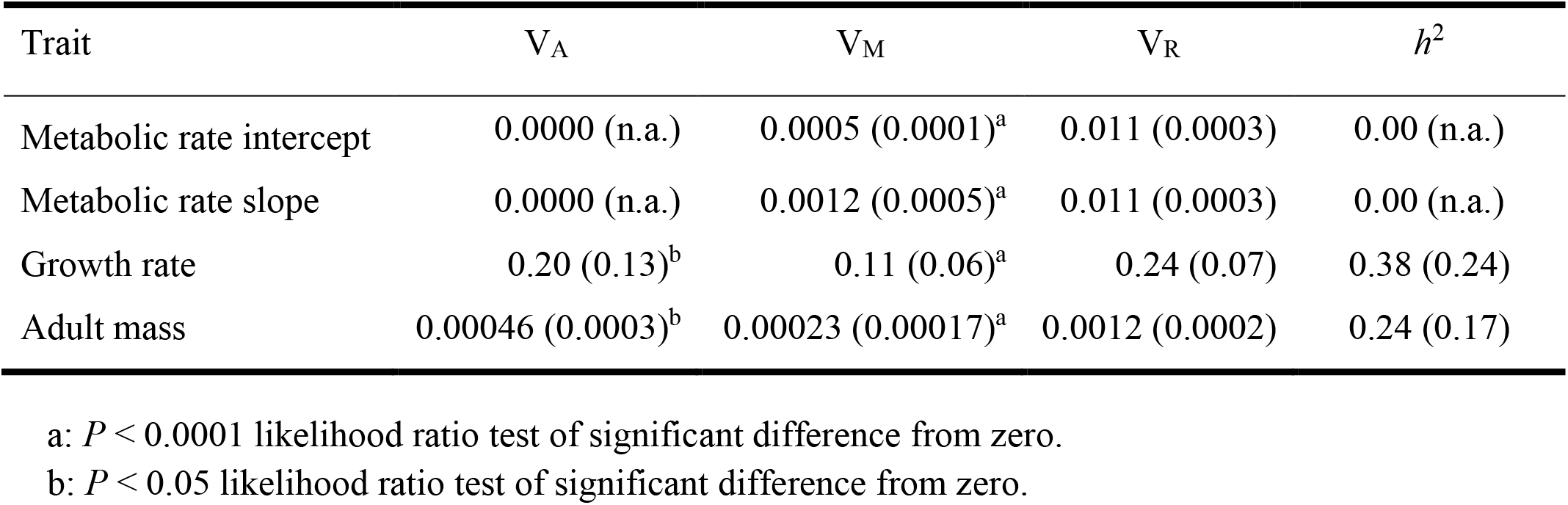
Additive genetic variance (V_A_), maternal variance (V_M_), phenotypic variance (V_P_) and heritability (*h*^2^) estimates (standard error shown) for ontogenetic metabolic scaling slope and intercept, growth rate and adult mass in *Nauphoeta cinerea* as estimated from univariate animal models. Significance levels of parameter estimates (one-tailed test; indicated by superscripts) were tested by likelihood ratio tests (Materials & Methods).

| Trait | $V_A$ | $V_M$ | $V_R$ | $h^2$ |
| --- | --- | --- | --- | --- |
| Metabolic rate intercept | 0.0000 (n.a.) | 0.0005 (0.0001)<sup>a</sup> | 0.011 (0.0003) | 0.00 (n.a.) |
| Metabolic rate slope | 0.0000 (n.a.) | 0.0012 (0.0005)<sup>a</sup> | 0.011 (0.0003) | 0.00 (n.a.) |
| Growth rate | 0.20 (0.13)<sup>b</sup> | 0.11 (0.06)<sup>a</sup> | 0.24 (0.07) | 0.38 (0.24) |
| Adult mass | 0.00046 (0.0003)<sup>b</sup> | 0.00023 (0.00017)<sup>a</sup> | 0.0012 (0.0002) | 0.24 (0.17) |

a: $P < 0.0001$ likelihood ratio test of significant difference from zero.
b: $P < 0.05$ likelihood ratio test of significant difference from zero.

**Figure 3.**
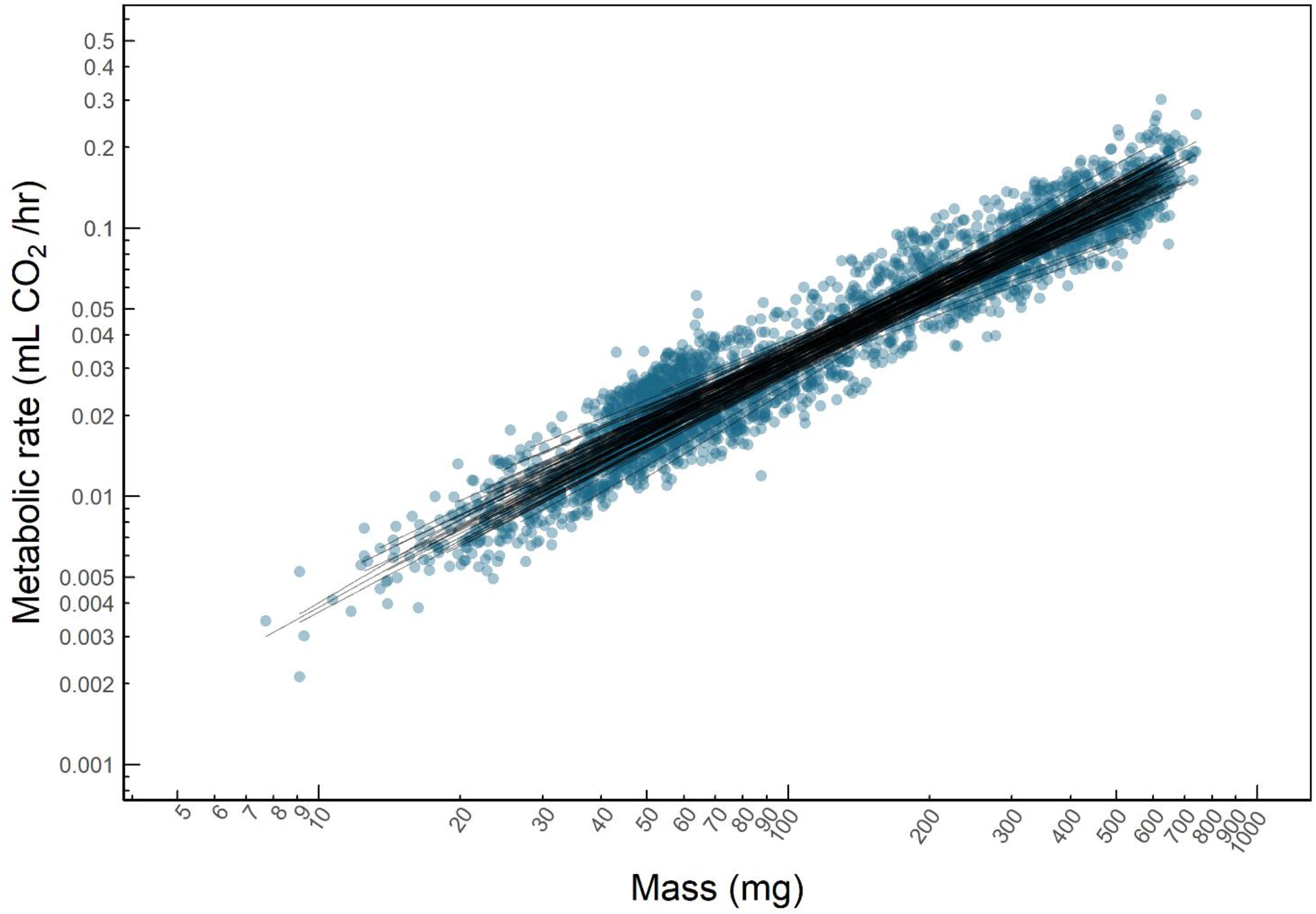
Metabolic rate and mass measurements repeated across ontogeny in speckled cockroaches *Naupoeta cinerea*. Points show individual-level paired measurements of mass and metabolic rate. Black lines represent dam-level (n = 134) linear regressions of log metabolic rate on log mass and visually depict the maternal effect variance in the ontogenetic metabolic scaling relationship. Axes show untransformed mass and metabolic rate values on a logarithmic (base 10) scale.

### Functional and selective sources of evolutionary constraint on metabolic allometry

Our observation of a lack of additive genetic variance in the metabolic scaling relationship is a novel result since there have been no previous attempts to estimate the narrow sense heritability of a metabolic scaling relationship. It has previously been shown that the ontogenetic metabolic scaling varies among individuals (Norin and Gamperl, 2018). A recent study in 10 clonal lines of *Daphnia magna* found that ontogenetic scaling slopes were responsive to temperature but found no evidence for broad sense genetic variance in scaling slopes despite the presence of broad sense genetic variance in the scaling intercept (Fossen et al., 2019). There are several reasons why the metabolic scaling relationship might exhibit such low additive genetic variance (Hoffmann and Blows, 2005). Two appealing hypotheses for the observed lack of heritability in the metabolic scaling relationship are: i) that there are functional constraints on the relationship between mass and metabolic (Kleiber, 1932, Schmidt-Nielson, 1984, West et al., 1997, West et al., 1999), and ii) that strong stabilising selection on combinations of mass and metabolic rate erodes genetic variance around the central tendency of the scaling relationship (Blows and Hoffmann, 2005, Hunt et al., 2007, Roff and Fairbairn, 2012, White et al., 2019). Previous studies have identified evidence for positive and negative correlational selection on combinations of mass and metabolic rate (Artacho et al., 2015) and on combinations of metabolic rates expressed at different ages (Pettersen et al., 2016), but there are too few estimates of selection on mass and metabolic rate to assess the pervasiveness of stabilising and directional selection on the covariation in mass and metabolic rate (Pettersen et al., 2018).

Importantly, functional and selective hypotheses for low genetic variance in the metabolic rate-mass relationship are not mutually exclusive. Rather, they are complementary since stabilising selection on mass and metabolic rate might arise due to underlying functional and developmental constraints on the covariation in mass and metabolic rate (Cheverud, 1988; Arnold, 1992). In future, it might be possible to reconcile the functional, genetic and selective sources of constraint on the evolution of metabolic scaling by investigating quantitative variation in the underlying mechanisms, such as the branching of vascular networks, and identify the genetic basis of those mechanistic variations and how they associate with variation in fitness. Recent notable studies have quantified variation in the branching architecture of vascular networks (Aitkenhead et al., 2020; Brummer et al., 2020 preprint) and explored how that variation corresponds to variation in metabolic scaling exponents (Brummer et al., 2017; Brummer et al., 2020 preprint). Yet it seems that at this stage it would require a prohibitively large phenotyping effort to link individual variation in underlying functional mechanisms to variation in fitness, which would be required to determine how variation in functional mechanisms are exposed to selection at the whole-organism level. What is more feasible now is to study how covariation in whole-organism mass and metabolic rate is associated with variation in fitness using high-throughput metabolic rate phenotyping (Kellermann et al., 2019, Videlier et al., 2019, Nagarajan-Radha et al., 2020). Such an approach would provide robust tests of hypotheses about the role of selection in the evolution of metabolic allometry.

We hypothesise that there is strong stabilising selection on combinations of mass and metabolic rate within populations, favouring a positive correlation between mass and metabolic rate with an allometric slope less than 1. This prediction could be tested by estimating lifetime reproductive success alongside paired measurements of mass and metabolic rate, allowing estimation of multivariate selection gradients for mass and metabolic rate (Lande and Arnold, 1983, Blows, 2007). We expect that the orientation of the multivariate fitness optimum (i.e. a fitness ridge on a three-dimensional fitness landscape) would be aligned in the same direction as the metabolic scaling relationship, with fitness rapidly declining as mass-independent metabolic rate deviates from the mean relationship. Another test of our hypothesis would be to use artificial selection on combinations of mass and metabolic rate to change the value of the metabolic scaling slope (following approaches for experimental evolution studies on morphological allometries by Egset et al., 2012, Bolstad et al., 2015 and Houle et al., 2019), after which artificial selection could be relaxed to see whether natural selection returns the scaling relationship to its original value. Such protocols have been used to demonstrate the evolvability of morphological allometries (Egset et al., 2012, Boldstad et al., 2015, Houle et al., 2019), but no studies have experimentally investigated the role of multivariate selection in shaping metabolic allometry.

## Materials & Methods

### Phylogenetically informed analysis of metabolic scaling

We performed an analysis of a large dataset of intraspecific metabolic scaling slopes from a range of vertebrate and invertebrate taxa compiled by Glazier (2005) (Glazier, 2005). The original (Glazier, 2005) dataset was for intraspecific metabolic scaling and included both ontogenetic and static scaling relationships. Typically, the mass ranges achieved over development are much larger (typically an order of magnitude or more) than the mass ranges among individuals of the same developmental stage (typically less than one order of magnitude). We therefore excluded static metabolic scaling slopes from the dataset by excluding all scaling slopes fit to mass ranges less than 0.9 orders of magnitude.

We used the package ‘rotl’ (Michonneau et al., 2016) in R and the Open Tree of Life (Hinchliff et al., 2015) database (https://ot39.opentreeoflife.org/about/open-tree-of-life) to generate a phylogenetic tree for the species in the Glazier (2005) dataset and to match the data to the tree. We then used ASReml-R (Butler, 2009) to run phylogenetic mixed models to test for phylogenetic signal in the ontogenetic metabolic scaling slopes. We also grouped the data based on whether species were endothermic or ectothermic, vertebrate or invertebrate, or aquatic, amphibious or terrestrial, and tested whether these groupings explained any of the variance in scaling slopes independently of shared history.

### Maintenance of outbred cockroach colonies

Approximately 2000 juvenile speckled cockroaches *N. cinerea* were purchased from an insect supplier (Brian’s Worms, Queensland, Australia) and maintained as outbreeding colonies in the laboratory. Colonies were maintained in four well-aerated plastic containers (40 L) with egg cartons provided for shelter and kept under ambient temperature and photoperiodic conditions. Cockroaches in the colonies were provided with ad libitum fresh carrot and dry dog biscuits (Optimum, crude protein 26%, crude fat 14%, energy density 355kcals/100g).

### Quantitative genetic breeding design

In the present study we used a quantitative genetic full-sibling, half-sibling breeding design in which phenotypic traits were measured in F_1_ offspring produced by mating F_0_ sires and dams sampled from the outbreeding colonies. In total, we measured the phenotypes of 1269 individual cockroaches in the F_1_ generation from 48 sires and 134 dams.

To form the F_0_ generation, two hundred late-stage juvenile cockroaches were removed from the colonies and allocated to individual containers. Juvenile F_0_ cockroaches were observed daily to check for individuals that had moulted to adult stage. Male and female *N. cinerea* become sexually receptive after 5 days from moult to adult, and female reproductive output and offspring quality remain relatively steady until about day 15, with reproductive output declining heavily from day 18 onwards (Moore and Moore, 2001, Moore and Sharma, 2005). Mature male and female cockroaches were paired for 24 hours between 5 and 14 days after the final moult to adult. Each sire was sequentially mated to between 2 and 4 dams. Sires were rested for at least 48 hours between mating events to reduce potential for sperm depletion. Dams were observed daily to check for newly hatched offspring. Clutch size ranged from 7 – 39 offspring, with a mean clutch size of 29.1. To form the F_1_ generation, ten offspring were haphazardly chosen from each clutch and maintained individually from hatch until maturity.

### Maintenance of experimental cockroaches

Cockroaches were maintained individually at 30.6 ± 0.98 (s.d.) °C and 74.0 ± 1.97 (s.d.) % relative humidity with a 12:12 hour light dark cycle. Individuals were housed within plastic tubes (50 mL) modified to allow air flow and to provide a dark shelter over half of the container. Containers were stored horizontally in racks kept inside humidity-controlled boxes (15 L) in a temperature-controlled room. Relative humidity inside the boxes was controlled by pumping air saturated at 28°C into the boxes, which were maintained at 30°C, thereby reducing the relative humidity to ~75% inside the boxes. Saturating air at 28°C was achieved by pumping air through water in Dreschel gas-washer bottles (Labtek, Queensland, Australia) kept in temperature-controlled incubators. Experimental cockroaches were provided with fresh carrot and dried fish pellets (crude protein 50%, crude fat 20%; crude fibre 4%; digestible energy 18.7 MJ/Kg; Ridley Aquafeeds, Queensland, Australia) on three days each week. Remaining food and faeces were removed the day after feeding and food was not replaced until the following day. The feeding regime meant that each cockroach had access to food for three 24 hour periods each week. The feeding rate used in the present study was determined through previous experiments to be sufficient to not limit juvenile growth and reproduction. At the start of the experiment all cockroaches were randomly assigned to blocks, and the location of blocks within the room were changed on a regular basis to minimise confounding effects of unknown factors such as temperature gradients in the room.

### Flow-through respirometry and mass measurements

We measured metabolic rate and mass repeatedly in individuals from early juvenile stages through to maturity. Mass of individuals was measured using a precision balance (Mettler Toledo, Ohio, USA) on the same day on which metabolic rate was estimated. Resting metabolic rate was estimated as the rate of carbon dioxide production (*V*CO_2_; mL h^−1^) in post-absorptive individuals at rest. *V*CO_2_ was measured using standard flow-through respirometry techniques (Schimpf et al., 2013). Briefly, individual cockroaches were placed in metabolic chambers made from plastic syringes (3 or 6 mL depending on the size of the cockroach). Air was drawn from outside and pumped through columns of soda lime and Drierite to remove CO_2_ and water vapour, respectively, then passed through metabolic chambers at a rate of 25 or 50 mL/min for 3 and 6 mL metabolic chambers, respectively. The concentration of CO_2_ in each airstream was measured using LI-840A gas analysers (LI-COR, Nebraska, USA), which recorded fractional concentrations of CO_2_ (parts per million, ppm) in the excurrent air at a frequency of 1 sample second^−1^. Flow rate was regulated to an appropriate flow rate for the chamber volume and cockroach size (25 – 50 mL min^−1^ STPD) using mass-flow controllers (Aalborg, Model GFC17, Orangeburg, NY). Flow rate was calibrated for each channel using a bubble flow meter (Bubble-O-Meter, OH). We calculated mean *V*CO_2_ during a 30 minute measurement period immediately following a 30 minute settling period (which was sufficient for cockroaches to reach a resting state), using the equation:

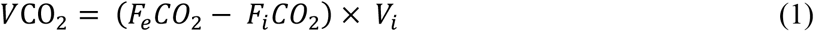

where *F*_*i*_*CO*_*2*_ is the fractional concentration of CO_2_ in the airstream incurrent to the chamber, *F*_*e*_*CO*_*2*_ is the fractional concentration of CO_2_ excurrent from the chamber, and *V*_*i*_ is the flow rate of air through the respirometry chamber in mL/min. *F*_*i*_*CO*_*2*_ was estimated by fitting a linear model to the baseline CO_2_ concentrations measured before and after a cockroach was placed inside the respirometry chamber. This model was used to interpolate the baseline CO_2_ concentration during the test period and to account for linear drift in the baseline. The interpolated baseline CO_2_ concentration was then subtracted from the CO_2_ concentration of the excurrent airstream during the 60 minute test period, from which mean *V*CO_2_ was calculated. Data traces were visually inspected for periods of CO_2_ production attributable to activity, which were subsequently removed from the period over which *V*CO_2_ was calculated. All measurements were made during the light phase of the day, which is the predominantly quiescent phase for the study species. Cockroaches were fasted for between 14 and 48 hours prior to measurement meaning they were in a post-absorptive state (Schimpf et al., 2012). Despite being post-absorptive, however, the cockroaches in the present study were still expending energy on growth, which was determined in previous experiments by measuring the growth rates of cockroaches fed at different frequencies (unpublished data). Thus, in the present study we have estimated resting metabolic rate, rather than standard metabolic rate. This is an important distinction because energy expended on growth contributes to variation in resting metabolic rate in juvenile organisms (Rosenfeld et al., 2015).

### Quantitative genetic analysis

We used linear mixed-effects models to estimate quantitative genetic parameters for the metabolic scaling slope and intercept, growth rate and adult mass. We used multivariate models to explore correlations among the traits in our model. We found no statistical support for a model that included correlations at the levels of additive genetic and permanent environment effects (χ^2^ = 1.2294, d.f. = 7, *P* = 0.76). There was support for a model that included trait correlations at the level of maternal effects (χ^2^ = 15.66, d.f. = 6, *P* = 0.0023), but these did not change our main findings, so in the interest of simplicity, we present the parameter estimates from our univariate models.

All statistical analyses were completed in R version 3.6.0 (R Core Team, 2017), using the ASReml-R package version 4.0 (Butler, 2009) and the package ‘plyr’ version 1.8.4 (Wickham, 2011). The general formulation of the linear mixed-effects models used was:

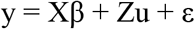

where β is a vector of fixed effects, X is an identify matrix for fixed effects, u is a vector of random effects, Z is an identity matrix for random effects, and ε is a residual error term, ε = *N*(0, σ^2^). We used restricted maximum likelihood to estimate model parameters. The allometric relationship between mass and metabolic rate is described by a power law on the measurement (untransformed) scale, which can be estimated using ordinary least squares regression to obtain the slope and intercept on a log-log scale. Hence, on a log-log scale, the regression slope and intercept represent the allometric exponent and a constant, respectively, on the untransformed scale (White and Kearney, 2014). Variances for the slope and intercept, and a slope-intercept correlation, were estimated using a random regression approach, with metabolic rate as the response variable and mass as a continuous predictor to model the mean slope and intercept of the scaling relationship. We modelled permanent environment (i.e. individual), additive genetic and maternal effect variances as random deviations around the mean slope and intercept using relatedness information from the pedigree (i.e. the animal model). We mean-centred log-transformed mass prior to model fitting, to remove the inherent covariance between the slope and intercept and help with model convergence. Other than mean-centring mass to model metabolic scaling slopes and intercepts, all other variables were left un-standardised in univariate models. Growth rate and adult mass were analysed in separate univariate models as single point estimates. To estimate growth rates, we fit linear regressions to the relationship between log mass and age for each individual. We then used the slope of the regression for each individual to calculate their growth rate as 100 × (10^slope^ - 1), which gives the daily growth increment as a percentage of mass at any given age. Cockroaches exhibit a sigmoidal pattern of growth on a logarithmic scale, so we only estimated growth rate from the linear portion of the growth curve, which represents maximal growth rate. Adult mass was measured when respirometry was performed after final moult to adult and was analysed on an untransformed scale in units of grams.

We compared nested models with log-likelihood ratio tests to test the statistical significance of our variance and correlation estimates. For each parameter, we used log-likelihood ratio tests to compare an unconstrained model to a model in which the focal variance estimate was constrained to very near zero (0.0001). In the case of metabolic scaling, the model simultaneously estimated the variance in the slope and intercept, and the slope-intercept correlation, of the linear relationship between log mass and log metabolic rate and each of these parameters were tested simultaneously and individually, with both approaches leading to same result. A permanent environment effect (i.e. repeatability) estimate was tested for metabolic scaling since metabolic rate was measured repeatedly in individuals. We found no statistical support for a permanent environment effects in the observed variation in the metabolic scaling relationship (χ^2^ = 0.16, d.f. = 3, *P* = 0.73). This was potentially influenced by the different number of records among individuals (most individuals had 5 – 6 records but some fewer than three records which may have reduced the precision of our estimate for individual-level effects). We retained the permanent environment effect in our final model of metabolic rate so as not to artificially inflate our estimates of additive genetic variance in the metabolic scaling slope.

We included sex and an experimental blocking effect as fixed effects in all models. For metabolic rate, we also included day of the week as a fixed effect to account for any possible effects on metabolic rate from the feeding schedule. For metabolic rate, the effect of day was significant (Wald test F _(5)_ = 185, *P* < 0.001) but the effects of sex (F_(2)_ = 5, *P* = 0.096) and block (F_(23)_ = 20, *P* = 0.64) were not significant for the metabolic scaling slope and intercept. For growth rate, the effects of sex (F_(2)_ = 14.8, *P* < 0.001) and block (F_(23)_ = 38.1, *P* < 0.05) were both significant. For adult mass, the effect of sex was significant (F_(2)_ = 611.8, *P* < 0.0001) but the effect of block was not significant (F_(23)_ = 26.9, *P* = 0.3).

## Acknowledgements

This study was supported by Australian Research Council grants to C.R.W., D.O-B., K.M. and M. D.H., and by an Australian Postgraduate Award to J.E.B. The authors would like to thank Lesley Alton, Pieter Arnold, Mark Blows, Candice Bywater, Evatt Chirgwin, Carmen da Silva, Katrina McGuigan, Adam Reddiex, Robbie Wilson and Hugh Winwood-Smith for helpful discussions, advice or practical help.

## Notes

### Competing Interest Statement

The authors have declared no competing interest.

